# Comparative Performance of Two Whole Genome Capture Methodologies on Ancient DNA Illumina Libraries

**DOI:** 10.1101/007419

**Authors:** María C. Ávila-Arcos, Marcela Sandoval-Velasco, Hannes Schroeder, Meredith L. Carpenter, Anna-Sapfo Malaspinas, Nathan Wales, Fernando Peñaloza, Carlos D. Bustamante, M. Thomas P. Gilbert

## Abstract

1. The application of whole genome capture (WGC) methods to ancient DNA (aDNA) promises to increase the efficiency of ancient genome sequencing.
2. We compared the performance of two recently developed WGC methods in enriching human aDNA within Illumina libraries built using both double-stranded (DSL) and single-stranded (SSL) build protocols. Although both methods effectively enriched aDNA, one consistently produced marginally better results, giving us the opportunity to further explore the parameters influencing WGC experiments.
3. Our results suggest that bait length has an important influence on library enrichment. Moreover, we show that WGC biases against the shorter molecules that are enriched in SSL preparation protocols. Therefore application of WGC to such samples is not recommended without future optimization. Lastly, we document the effect of WGC on other features including clonality, GC composition and repetitive DNA content of captured libraries.
4. Our findings provide insights for researchers planning to perform WGC on aDNA, and suggest future tests and optimization to improve WGC efficiency.

## INTRODUCTION

The introduction of next-generation sequencing (NGS) marked a dramatic turning point in aDNA investigation, enabling the study of genome scale datasets (e.g. Green *et al.* 2010, Meyer *et al.* 2012, Orlando *et al.* 2013, Rasmussen *et al.* 2014). Nevertheless, the DNA quality within most archaeological samples continues to hamper the field’s development. Specifically, the fragmented and damaged nature of aDNA molecules, coupled with high levels of exogenous contaminant DNA, have required investment of significant resources in order to enable generation of meaningful levels of sequence data (Knapp & Hofreiter 2014). In response to this challenge, several key methodological improvements have been developed. These include optimization of DNA extraction and library preparation protocols to better retain and incorporate damaged endogenous DNA (e.g. Dabney *et al.* 2013, Gansauge & Meyer 2013, respectively), and the introduction of large-scale hybridization-capture based targeted enrichment techniques optimized for aDNA (e.g. Briggs *et al.*, 2009, Burbano *et al.*, 2010, Maricic *et al.* 2010, Schuenemann et al, 2011, Ávila-Arcos et al, 2011, Fu *et al.* 2013).

Until recently, targeted enrichment approaches were limited to relatively small fractions of predefined regions of an already characterized genome, (e.g. Maricic *et al.* 2010, Schuenemann *et al.* 2013, Fu *et al.* 2013). However, a method introduced by Carpenter *et al.* (2013), termed Whole-genome In-Solution Capture (WISC), showed considerable genome-wide enrichment of human aDNA sequencing libraries with very low initial concentrations of endogenous DNA. In parallel, a commercial method, MYbaits-WGE (MYbaits Whole Genome Enrichment, henceforth referred to as “MYbaits”), was developed by the company MYcroarray (Ann Arbor, MI, USA) following a similar methodological principle (Gnirke *et al.* 2009) but with some subtle differences in the protocol (for an application of the method see Enk *et al.*, 2014). In broad terms, the approach involves the construction of an RNA ‘bait’ library by transcribing modern genomic DNA that has been fragmented and ligated to T7 RNA polymerase promoters. The synthesis of RNA from the T7-ligated modern reference genomic DNA is carried out using biotinylated UTPs, and the RNA products are then hybridized in solution to aDNA libraries. The hybridized fragments can then be retrieved using streptavidin-coated magnetic beads, while the unbound molecules are washed away. Captured library molecules can subsequently be amplified and sequenced (Carpenter *et al.* 2013, Enk *et al.* 2014).

Although the fundamental molecular principle of both methods is similar, it is unclear if they performed equally well, and as such, whether there is a user benefit in employing one over the other. To explore this issue, and to better understand the parameters affecting the success of capture experiments, we enriched and sequenced a series of ancient human DNA libraries using both WISC and MYbaits. Using the data, we describe the effect that different bait lengths and hybridization times have on the resulting fold enrichment, when applying each protocol to aDNA libraries with variable levels of initial endogenous DNA content. To a lesser extent we also explore the potential role of differences in hybridization times. Furthermore, we investigate the performance of both whole genome capture (WGC) methods on single-stranded libraries (SSL), which have been previously shown to contain higher levels of endogenous DNA than standard double-stranded libraries (DSL) (Meyer *et al.*, 2012, Gansauge & Meyer, 2013). We examined the effect on fold enrichment of applying WGC methods to this particular type of libraries.

## MATERIALS AND METHODS

### DNA extraction and aDNA library preparation

We generated sequence data from DNA extracted from eight archeological human skeletal samples originating from a range of different archaeological contexts and environmental conditions, dated to between 300 and 2500 years BP (Table 1). We deliberately chose samples from different contexts and with variable amounts of endogenous DNA as determined through shotgun sequencing (0.2-8.0%) to assess the performance of the two different WGC methods on samples of different quality. All DNA extraction and library preparation steps were performed in dedicated clean laboratories at the Centre for GeoGenetics in Copenhagen, Denmark, to prevent contamination with modern DNA. Before extraction, the tooth samples were cleaned with 10% bleach solution and then UV-irradiated for 2 min on each side to cross-link potentially contaminant DNA to the surface. Part of the tooth root was then excised and the inside of the tooth was drilled to produce approximately 200 mg of powder. The powder was digested in 5 ml of an EDTA-based digestion buffer containing 0.25 mg/mL Proteinase K. DNA was then isolated using a silica-based extraction method (Rohland and Hofreiter 2007). Samples were eluted in 60 μl TET buffer.

**Table 1.**
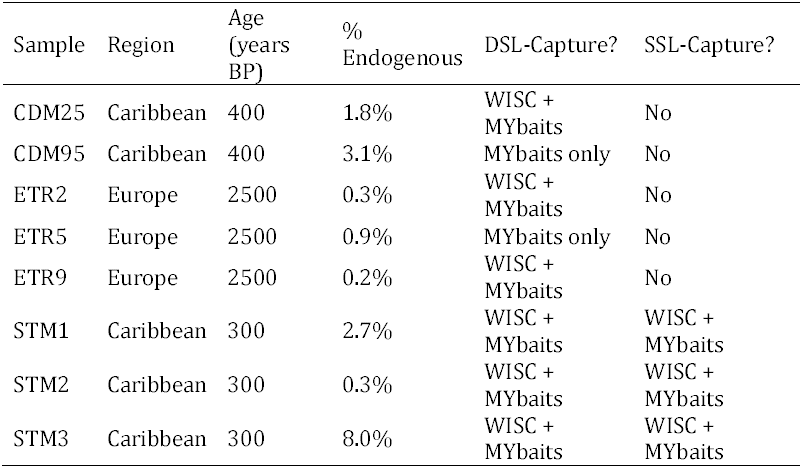
Description of samples and WGC scheme used for each. Provenance and approximate age of the samples are shown, along with the pre-capture endogenous content of the DSL libraries built from them. The type of library (DSL: Double-stranded library; SSL: Single-stranded library) built and the whole genome capture method(s) used are depicted in the last two columns.

| Sample | Region | Age<br>(years<br>BP) | %<br>Endogenous | DSL-Capture? | SSL-Capture? |
| --- | --- | --- | --- | --- | --- |
| CDM25 | Caribbean | 400 | 1.8% | WISC +<br>MYbaits | No |
| CDM95 | Caribbean | 400 | 3.1% | MYbaits only | No |
| ETR2 | Europe | 2500 | 0.3% | WISC +<br>MYbaits | No |
| ETR5 | Europe | 2500 | 0.9% | MYbaits only | No |
| ETR9 | Europe | 2500 | 0.2% | WISC +<br>MYbaits | No |
| STM1 | Caribbean | 300 | 2.7% | WISC +<br>MYbaits | WISC +<br>MYbaits |
| STM2 | Caribbean | 300 | 0.3% | WISC +<br>MYbaits | WISC +<br>MYbaits |
| STM3 | Caribbean | 300 | 8.0% | WISC +<br>MYbaits | WISC +<br>MYbaits |

Following extraction, the DNA was divided into two aliquots of 30 μl. Each of these were built into Illumina libraries, using a double- and single stranded protocol, respectively. The single-stranded libraries were built following a previously published protocol (Gansauge & Meyer, 2013) but without first removing deoxyuracils. The double-stranded libraries were prepared using a blunt-end library preparation kit from NEB (E6070) and blunt-end modified Illumina adapters (Meyer and Kircher, 2010). The libraries were prepared according to the manufacturer’s instructions, although with minor modifications as outlined below. The initial nebulization step was skipped because of the fragmented nature of aDNA. End-repair was performed in 50 μl reactions with 30 μl of DNA extract. The end-repair cocktail was incubated for 20 min at 12°C and 15 min at 37°C and purified using Qiagen MinElute silica spin columns following manufacturer’s instructions and eluted in 30 μl. After end-repair, Illumina-specific adapters (Meyer and Kircher, 2010) were ligated to the end-repaired DNA in 50 μl reactions. The reaction was incubated for 15 min at 20°C and purified using Qiagen QiaQuick columns before being eluted in 30 μl EB. The adapter fill-in reaction was performed in a final volume of 50 μl and incubated for 20 min at 37°C followed by 20 min at 80°C to inactivate the *Bst* enzyme. Libraries were then amplified and indexed in a 50 μl PCR reaction, using 15 μl of library template, 25 μl of a 2x KAPA U+ master mix, 5,5 μl H2O, 1,5 μl DMSO, 1 μl BSA (20 mg/ml), and 1 μl each of a forward and reverse indexing primer (10 μM). Thermocycling conditions were 5 min at 98°C, followed by 10-12 cycles of 15 sec at 98°C, 20 sec at 60°C, and 20 sec at 72°C, and a final 1 min elongation step at 72°C. The amplified libraries were then purified using Agencourt AMPure XP beads and eluted in 30 μl EB. Between 2-6 μl of the indexed DNA libraries were then quantified on an Agilent Bioanalyzer, pooled in equimolar amounts, and sequenced together with other samples on a HiSeq 2000 lane.

### Whole Genome Capture

The remaining fraction of the libraries was subdivided, and re-amplified for 8-12 cycles using primers IS5 and IS6 (from Meyer & Kircher, 2010) and the same PCR conditions as above to obtain a minimum of 100 ng of library template. The libraries were then captured using the two different WGC methods. WISC, which makes use of homemade RNA probes, was carried out as described in Carpenter *et al.* (2013), with a bait-library hybridization time of 66 hours. For the MYbaits capture, a human whole genome enrichment kit MYbaits-HuWGE (MYcroarray, Ann Arbor), which relies on pre-made RNA probes and adapter blockers built using proprietary protocols, was used following manufacturer’s instructions with 24 hours for bait-library hybridization. Captured libraries were re-amplified with IS5 and IS6 for 10-20 cycles (depending on library and capture method, see Table 2 for total number of PCR cycles per sample) and using the same PCR conditions as above. Subsequently, the libraries were purified, quantified and pooled as described above, and submitted for sequencing on the HiSeq 2000.

**Table 2.**
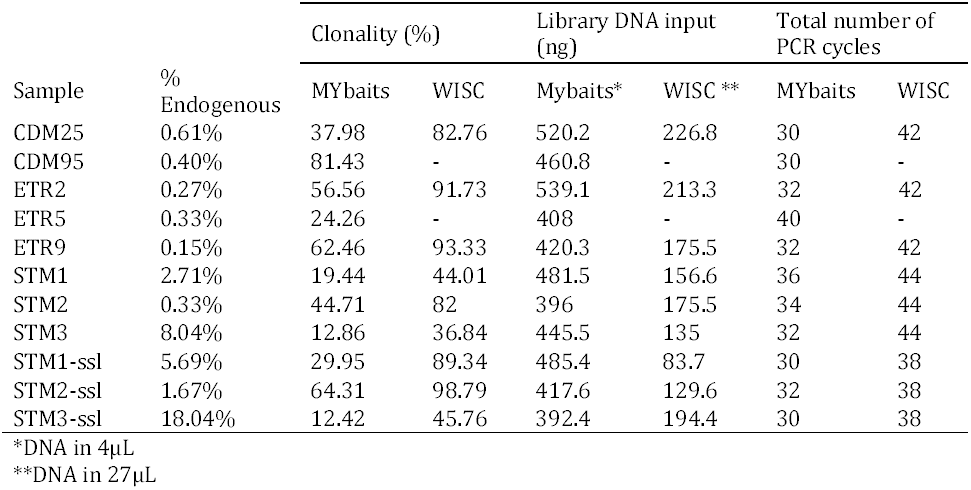
Lower pre-capture endogenous content results in higher clonality (measured as the fraction of the total mapped reads that are clonal (PCR) duplicates) in post-capture libraries. The total number of cycles includes two pre-capture and one post-capture amplification rounds. Higher clonality in WISC experiments is likely caused by the higher number of total PCR cycles used in these experiments, which in turn is due to less DNA used as input of the capture experiment. This correlation follows a logarithmic curve as illustrated in Supplementary Fig. S7.

### Fastq filtering and mapping

Libraries were sequenced on an Illumina HiSeq2000 in 100-cycle single-end runs. The base calls were performed using the Illumina software CASAVA 1.8.2 and the output fastq files were processed with the following software and scripts. First, reads were filtered with AdapterRemoval (Lindgreen 2012) by trimming adapter sequences as well as N’s and low quality stretches and discarding fragments shorter than 30 bp. Reads passing filters were then mapped using BWA Version: 0.7.5a-r405 (Li & Durbin 2009) to the hg18 reference genome, setting the minimum mapping quality to 25. Next, Samtools rmdup (Li *et al.* 2009) was used to remove PCR clones while reads mapping to more than one unique region in the genome were also excluded by controlling for the XT, XA and X1 tags in the bam alignment. Fragmentation and damage patterns were calculated and plotted using MapDamage2 (Ginolhac *et al.* 2011, Jonsson *et al.* 2013).

### Contamination estimates

To estimate the contamination fraction for each experiment, we mapped the data to the whole nuclear genome (hg18) as well as to a consensus mitochondria (mt) sequence that was generated separately for each sample [STM1, STM2, STM3 (Schroeder et al, unpublished data)]. We only retained reads that mapped to the consensus mt with a mapQ > 30, and excluded reads with potential alternative mapping coordinates to the nuclear genome by controlling for XT, XA and X0 tags in the bamfile. This has the effect of reducing the number of reads that map both to the mitochondrial genome and to nuclear copies of mitochondrial genes (numts). We then used a method detailed in the Supplemental Information section 5 of (Fu *et al.* 2013) that generates a moment-based estimate of the error rate and a Bayesian-based estimate of the posterior probability of the contamination fraction. We ran three chains of 50,000 iterations for the Monte Carlo Markov Chain and discarded the first 10,000, as was done in (Fu *et al.* 2013). We assessed convergence of the chain by visualizing the potential scale reduction factor (PSRF) and verifying that the median of PSRF is below 1.01 for all cases (Gelman & Rubin 1992, Plummer *et al.* 2006).

### Length distribution, repeat and GC content analysis

Bam files were intersected with the hg18 UCSC 50mer mapability tracks (downloaded from ftp://hgdownload.cse.ucsc.edu/gbdb/hg18/bbi/crgMapabilityAlign50mer.bw) using bedtools intersect (version 2.15.0, https://github.com/arq5x/bedtools) and requiring at least 50% of the read overlapping with a region with a mapability value of 0.5 or less. Unix scripts were used to calculate the GC content fractions and length distribution. Plots were generated using Rstudio (http://www.rstudio.org/).

## RESULTS

Shotgun sequencing of eight ancient human DNA libraries revealed pre-capture endogenous content ranging between 0.2% and 8% (Table 1). Prior to sequencing, human genomic DNA was enriched for two of the libraries using the MYbaits protocol and for six of the libraries using both the WISC and MYbaits protocols. Additionally, for three of the samples (samples STM1-3), SSL were built from the same DNA extract used to build the DSL. These three SSL were shotgun sequenced and also subjected to both capture schemes and sequenced on the HiSeq 2000 (Table 1). Sequenced reads that mapped to the human genome reference showed the damage and fragmentation patterns characteristic of aDNA (Briggs *et al.* 2007) and were consistent with the type of library build method used to generate them (Supplementary figures S1-4).

### Enrichment rates on DSL

To evaluate the performance of capture experiments, we computed the percentage of reads in each of the sequenced libraries that matched the human genome reference sequence, as well as the percentage of non-clonal, uniquely mapped reads that matched the reference. To estimate the fold enrichment for each sample, we compared the unique endogenous fraction of the captured libraries with that of the shotgun libraries (Ávila-Arcos *et al.*, 2011).

Regardless of the capture method, an increase of the endogenous content was consistently observed in all captured DSL libraries when compared to their pre-capture counterpart (Table 3). Informative rates of enrichment ranged between 1.9 and 8.6 fold. Although the sequences generated from the WISC libraries initially exhibited a higher fraction of human DNA, this derived from higher levels of clonal reads (Table 2), and the actual informative (unique) rate of enrichment was consistently higher for MYbaits experiments. Fold enrichment values ranged between 2.9 and 8.6 (mean 5.2) for MYbaits libraries and between 1.9 and 5.6 (mean 3.6) for WISC (Table 3), consistent with reported values (2 to 13 fold) in Carpenter *et al.* (2013).

**Table 3.** Fold enrichment of WGC by method and type of captured library. Numbers represent the unique informative (non-clonal) enrichment of post-capture libraries when compared to their pre-capture counterpart.

| Sample | DSL |  |  | SSL |  |  |  |
| --- | --- | --- | --- | --- | --- | --- | --- |
|  | % End<sup>a</sup> | WISC | MYbaits | % End<sup>b</sup> | shotgun<br>SSL<sup>c</sup> | WISC<sup>c</sup> | MYbaits<sup>c</sup> |
| CDM25 | 0.6% | 3.04 | 4.83 | - | - | - | - |
| CDM95 | 0.4% | - | 5.906 | - | - | - | - |
| ETR2 | 0.3% | 2.70 | 5.10 | - | - | - | - |
| ETR5 | 0.3% | - | 2.90 | - | - | - | - |
| ETR9 | 0.2% | 1.93 | 4.13 | - | - | - | - |
| STM1 | 2.7% | 5.53 | 5.96 | 5.7% | 2.10 | 0.42 | 2.22 |
| STM2 | 0.3% | 5.66 | 8.62 | 1.67% | 5.14 | 0.12 | 1.52 |
| STM3 | 8.0% | 2.74 | 3.73 | 18.07% | 2.24 | 1.15 | 1.68 |

<sup>a</sup>Endogenous DNA content of pre-capture DSL
<sup>b</sup>Endogenous DNA content of pre-capture SSL
<sup>c</sup> Enrichment values are in relation to pre-capture DSL

In contrast to previous observations (Carpenter *et al.* 2013; Enk *et al.* 2013) that lower initial endogenous DNA concentrations result in higher enrichment, we did not observe a clear dependence of enrichment rates on initial endogenous values. For example, among the pre-capture libraries, unique endogenous content values ranged between 0.2% and 8.0%, and for these extreme values the fold enrichment was very similar (4.1 fold and 3.7 fold, respectively, Table 3). This discrepancy could also be attributed to the small size of our dataset.

### Bait length influences the read length distribution of captured reads

The endogenous DNA sequences generated from captured libraries were consistently longer than those in shotgun libraries (Fig. 1a and 1b and Supplementary Figs. S5-6), in agreement with observations in previous WGC studies (Carpenter *et al.* 2014; Enk *et al.* 2014). Furthermore, libraries captured with WISC showed stronger bias against short fragments, and consequently a higher proportion of longer reads, than those enriched with MYbaits. To investigate if any physical properties of the baits could explain this discrepancy, we plotted the distribution of both bait libraries (see Materials and Methods) and observed the distribution of WISC probes was wider than MYbaits, with a higher fraction of the former exceeding 500 bp (Fig. 1c), whereas MYbaits had a narrower distribution between roughly 100bp and 500bp.

**Fig. 1.**
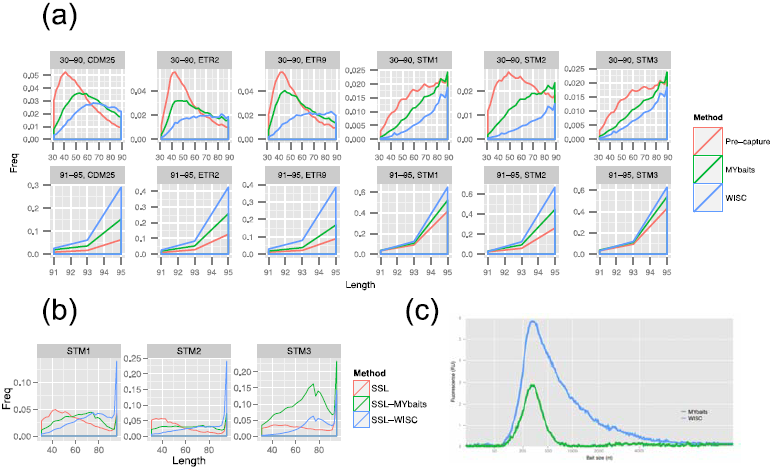
WGC preferentially retrieves longer fragments in sequencing libraries. The read length distribution of pre-capture and post-capture libraries is shown for (a) double-stranded libraries (DSL) and (b) single-stranded-libraries. In (a) the x axis is split in <90 bp and >90 bp to adjust the scale and better illustrate the higher concentration of short reads in the pre-capture libraries (pink line) and the bias observed against these in capture experiments (green and blue lines) where longer fragments are preferentially retrieved. (b) Illustrates that the relative gain of shorter fragments obtained by building a SSL, is lost by capturing these types of libraries. The plot shown in (c) depicts the bioanalyzer profile of the bait libraries revealing that for WISC a wider tail is observed for longer baits, which might explain the stronger bias in favor of longer fragments by this particular method.

### Effect of hybridization times

Although one might intuitively expect a correlation between incubation time and hybridization success, we observed no such behavior. The time of hybridization in WISC experiments was 66 hours, versus 24 hours for MYbaits. Our results suggest that the increased time utilized in WISC did not have a clear positive effect in terms of informative enrichment; however, we did not directly compare hybridization times between experiments performed with the same protocol (i.e., WISC for 24 hrs. vs. 66 hrs.).

### Clonality

The effect of capture on levels of sequence clonality was consistent with previous reports (Ávila-Arcos *et al.*, 2011). Firstly, sequence clonality (measured as the fraction of the total mapped reads that are clonal duplicates) in captured libraries was consistently higher than in pre-capture libraries. Secondly, the lower the endogenous content in the shotgun library, the higher the clonality in the post-capture ones. Furthermore, our results showed a logarithmic decrease of clonality with initial endogenous content (Table 3, Supplementary Fig. S7). In order to achieve the amounts of DNA required for WGC (both protocols requiring the same initial 100-500 ng), all libraries required two rounds of amplification prior to capture, and a further round of amplification post capture (see Materials and Methods). In fact, the amount of library DNA used as input for WISC was lower than that used for MYbaits (Table 2), which made necessary additional PCR cycles for the former in order to reach optimal concentrations for sequencing. As a result, WISC libraries consistently exhibited higher levels of clonal reads than MYbaits (Table 2).

### Whole Genome Capture on SSL

An unexpected observation was that, despite the SSL having higher endogenous DNA contents prior to capture than the DSL, both capture methods performed poorly on the SSL, with endogenous contents increasing only marginally and even decreasing relative to pre-capture values (Table 2). This poor performance of both WGC methods on SSL can be explained by the fact that the biggest gain in SSL is due to the recovery of short DNA fragments, which are then lost when WGC is performed with the parameters assayed herein. Further optimization of these parameters (e.g. hybridization temperature), could in principle overcome this limitation (see Discussion).

### Contamination

An important consideration when working with human aDNA is to control for potential modern human contamination. Reads derived from such contaminating DNA would intuitively be expected to have a longer average lengths than aDNA reads; hence, we investigated whether the fact that capture biased toward longer DNA molecules, resulted in higher contamination values in the captured libraries. We estimated the mitochondria (mt) contamination values for the three samples for which both DSL and SSL were subjected to WGC (see Material and Methods). We observed less mt contamination in the post capture libraries, suggesting the longer reads for those samples do not derive from a contaminant source. Although our sample size is small, it is an encouraging result for the use of capture methods in aDNA (Supplementary Table 1).

### GC content in WGC libraries

Previous capture studies have reported differences in GC content between pre- and post-capture libraries (Gnirke *et al.* 2009, Carpenter *et al.* 2013). We explored this factor in our dataset by measuring the GC fraction in pre- and post-capture reads and studying their distribution per experiment. We did not observe a specific pattern of increase or decrease of average GC contents in post-capture libraries, but instead a narrowing of the GC distribution in DSL libraries (Fig. 2a and Supplementary Table 2). Average GC content ranged between 42-49% in pre-capture DSL and between 44-46% and 43-45% in post-capture WISC and MYbaits libraries, respectively (Supplementary Table 2). Interestingly we noticed that the average GC of pre-capture SSL (35-38%) was lower that the than DSL, and that the SSL GC distributions were shifted downwards, as shown in the histograms in Fig. 2c. Remarkably, the GC content increased again when SSL were captured (Figs. 2b-c and Supplementary Table 2). Post-capture SSL mean GC contents ranged between 42-45% and 40-42% for WISC and MYbaits, respectively. We hypothesize that this bias towards specific GC values in post-capture libraries, relates to the particular selection of the hybridization temperatures and times in capture experiments.

**Fig. 2.**
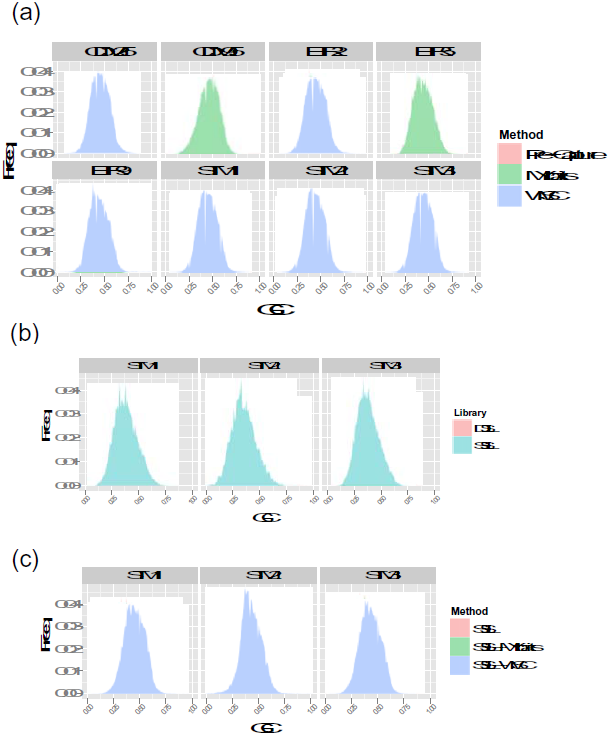
GC content distribution of WGC libraries is narrower than pre-capture ones. GC distributions in DSL libraries (a) show a subtle narrowing in the post-capture experiments (See Supplementary table 2 for exact ranges and summary statistics). Whereas the GC distribution in pre-capture SSL libraries displays a shift toward lower GC values when compared to pre-capture DSL (b), these values increase in post-capture SSL (c).

### Repeat enrichment in WGC libraries

Despite the inclusion of blocking agents (human Cot-1 and salmon sperm DNA) in both WGC protocols to “mask” repetitive elements in the library and avoid bait hybridization to these, a higher proportion of reads in post-capture libraries mapped to repetitive regions in the genome (Fig. 3). In particular, we observed that libraries captured with WISC displayed a higher fraction of repeats. To investigate if this was tied to the biased selection of longer reads, we computed and plotted the length distribution of reads within and outside repeats for each type of experiment (Supplementary table 3). These distributions confirmed that reads within repeats are on average longer in the pre-capture libraries, which probably drives these to preferential capture over the non-repeat ones, as evidenced by an also longer average length of reads within repeats in both post-capture library types (Supplementary Table 3).

**Fig. 3.**
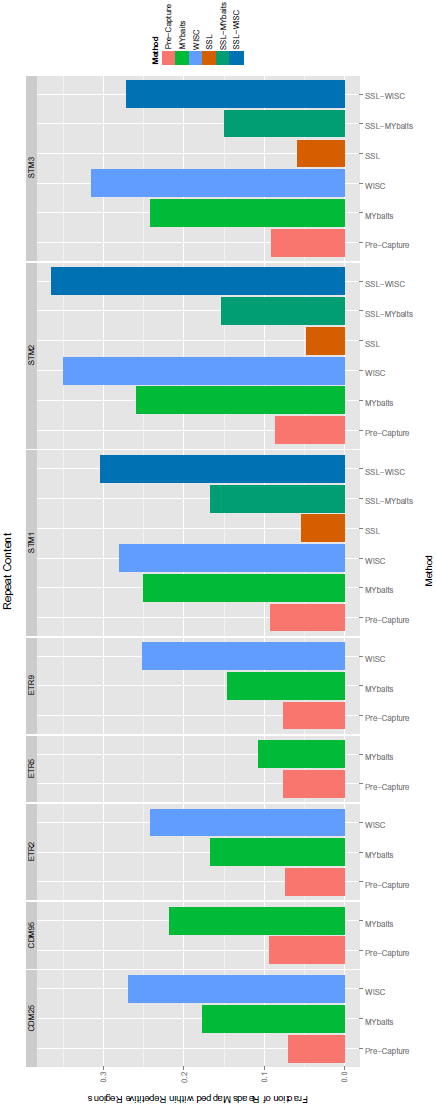
Fraction of reads mapping to repeated regions in the human genome (see Materials and Methods) in pre- and post-capture libraries. Post-capture libraries display a higher fraction of reads within repeats than pre-capture. This pattern is likely tied to the preferential capture of longer reads, which are also enriched in repeats in pre-captured libraries (Supplementary Table 3)

## DISCUSSION

By comparing the performance of WISC and MYbaits in enriching for endogenous human DNA in ancient DNA extracts, we have been able to pinpoint potential factors influencing the dynamics of WGC experiments. The assessment of the subtle differences between both approaches to in-WGC enables us to draw insights on two variables that may the affect capture efficiency – bait length distribution and hybridization time. Our data furthermore provides insights into the effect of blocking agents, and first insights into the performance of whole-genome enrichment methods on SSL.

Although the experimental design and parameters used in this study seem to suggest an apparent benefit of one of the methods over the other, we strongly caution that batch effects could be playing an important role under these settings, hence discourage such interpretation from our results. Likewise, it is worth considering that even though there is a certain convenience in using a pre-made kit (MYbaits), our observations point to specific factors that can be optimized in the in-house method (WISC) namely bait length distribution and hybridization parameters. Knowing the relevance of such parameters in WGC, gives users the flexibility of customizing their capture experiment to match the particularities of each aDNA library (see below).

### Role of bait length distribution on the efficiency of WGC

Bait length distributions (Fig. 1c) differ mainly in that WISC shows a wider range and longer bait lengths. This in principle could account for the marked retrieval of longer reads in the WISC compared with the MYbaits experiments (Figs. 1a-b). Following this rationale, the higher success of the latter could be explained by its ability to better access a fraction of the sequencing library, specifically that with the smaller fragments, while this fraction remains inaccessible due to the higher concentration of longer baits used in WISC. An important consequence of this feature was the poor and even unsuccessful outcome of capturing SSL, which include a higher fraction of short fragments, with either method. At the same time, this limitation reveals an important area for future development in the context of WGC experiments.

Although there was a small, yet consistent, benefit in the MYbaits over the WISC, it would be rash to conclude that the MYbaits method always outperforms WISC. These results were generated using a single batch of WISC bait versus a single batch of MYbaits. Given that (i) bait lengths will likely vary between batches as a result of initial template DNA fragmentation, and (ii) our hypothesis that bait length may play a key role in retaining shorter DNA fragments, we believe it is more than likely our results simply reflect the fact that in these batches tested the WISC bait were slightly longer than the MYbaits. Future studies that examine the role of bait length in capture success will be needed to further examine this hypothesis.

### Preferential retrieval of repeats in WGC experiments

Despite the inclusion of blocking agents, both capture methods resulted in a higher proportion of reads mapping to repetitive regions in post-capture compared to the pre-capture libraries. This enrichment of repeats in capture experiments has also been observed previously in WGC experiments, despite the inclusion of organism-specific Cot-1 DNA in the capture reactions (Carpenter *et al.* 2013 and Enk *et al.* 2014). Some of the possible explanations, as described by Enk *et al.* (2014), include more rapid association rates for repetitive sequences and jumping PCR in post-capture amplification. Furthermore, it has been observed that, against intuition, in some cases Cot-1 can enhance non-specific hybridization and hybridization to conserved repetitive elements (Newkirk *et al.* 2005). Also, batch-to-batch differences in the ability of commercial Cot-1 preparations to reduce hybridization to repetitive sequences have been reported (Carter *et al.* 2002). Our analyses show that another element to consider is the length of repetitive fragments, which we found to be longer than non-repetitive fragments and probably preferentially captured. In summary, considerations in this regard, along with further tests on the amount of this type of blockers, are needed in order to increase our understanding of their effects and to improve the capture efficiency.

### Cost-benefit of WGC experiments

We have demonstrated that WGC, at least for the two currently available methods, is an effective way to enrich endogenous content in aDNA sequencing libraries. However, it is important to consider that despite the success of its application, WGC would only represent a practical and cost-effective advantage for whole genome sequencing from ancient samples when applied on libraries with endogenous contents above 1%. Assuming an average read length of 60 bp, approximately 200 lanes worth of sequencing on the HiSeq2000 would be required to reach 1x of the human genome from a library with 1% endogenous (assuming high complexity in the library); however, only 23 would be needed if the sample is enriched to 8.6 fold, which is the maximum enrichment value observed in this study. Although the cost of sequencing 23 lanes is still very high, depending on the research question, less than 1x genome coverage could be enough to provide valuable information about the sample (e.g. Skoglund *et al.* 2012, Sánchez-Quinto *et al.*, 2012, Carpenter *et al.* 2013) Therefore, samples with less than 1% endogenous could in principle also be considered for WGC when the research question does not require information from the complete genome. Ultimately, optimization of capture enrichment protocols might enhance the efficiency and lower costs enough to make these samples with very little endogenous DNA accessible to genomic scale investigations.

With these observations in hand, we believe that an a *priori* knowledge of the endogenous content of a sample is crucial before deciding the enrichment strategy, or if it is even worth the investment of a WGC reaction. An initial shotgun screening can provide this information. Even with as few as 1 million reads (1/250 of a HiSeq2000 lane), characteristics of the library such as complexity, read length distribution and endogenous content can be retrieved and inform the selection of the strategy to follow, whether it is WGC (for reads with ∼1% endogenous, or even less when just a fraction of the genome is informative, and a higher proportion of long reads) or SSL (for very fragmented libraries). In fact, both strategies can be complementary and applied in parallel (but not consecutively) to cover a wider spectrum of read lengths. Additional optimization of WGC bait lengths and hybridization parameters might eventually overcome this limitation and make accessible the shorter fragments to WGC approaches.

In summary, we believe that our observations, though based on a small dataset and more descriptive than exhaustive, can be of value for researchers planning enrichment experiments. As new methods in aDNA are being developed at an unprecedented pace, being able to discern the best approach for a given sample is of the utmost relevance in order to take advantage of the methods and avoid wasting resources and precious sample material. New extraction methods, library preparation, sequencing and enrichment protocols are in their infancy in aDNA but promise to unlock fascinating information from ancient samples at an unparalleled scale. Consequently, it is important to have standards and guidelines to choose the best approach for any given project.

## Acknowledgements

The authors thank the staff at the Danish National High throughput Sequencing Centre for Technical Assistance, Jay Haviser and Corinne Hofman for providing samples, Jean-Marie Rouillard and Jake Enk for thoughtful input into the manuscript, and Alexandra Adams for technical support during WISC experiments. The authors acknowledge the following grants for generous financial support: Danish Research Foundation grant DNRF94, Danish Council for Independent Research grant 10-081390, Lundbeck Foundation grants R52-A5062 and Marie Curie Actions grant EUROTAST FP7-PEOPLE-2010, George Rosenkranz Prize for Health Care Research in Developing Countries, National Science Foundation award DMS-1201234, NIH NRSA postdoctoral fellowship (grant no. 5 F32 HG007342), Swiss National Science Foundation and NEXUS1492 (ERC Synergy Project Grant Agreement n° 319209)

## CONFLICT OF INTEREST

The authors declare no conflict of interest.

